# Greater Expression of DNA Repair Pathways in Sharks vs. Rays/Skates Based on Transcriptomic Analyses

**DOI:** 10.1101/2025.04.20.649719

**Authors:** Colton R. Simmons, Schaefer L. Grant, Lucia Llorente Ruiz, David W. Kerstetter, Manasi Pimpley, Jean J. Latimer

**Author notes:** Permanent address: 3200 S University Dr, Fort Lauderdale, FL 33328, United States.

## Abstract

Elasmobranchs are an understudied taxon of cartilaginous fishes. DNA repair studies have been performed in very few elasmobranchs. Because DNA repair maintains the integrity of the genetic code, it is important for the survival of elasmobranchs in increasingly polluted oceans. Oil spills, for example, have been shown to cause DNA adducts in marine animals. We hypothesized that four elasmobranch species would show differential DNA repair expression. Dermal tissue was harvested from nurse sharks (*Ginglymostoma cirratum*), spiny dogfish (also considered to be sharks) (*Squalus acanthias*), yellow stingrays (*Urobatis jamaicensis*), little skates (*Leucoraja erinacea*), and RNA was isolated. RNA sequencing was performed using the holocephalan Australian ghostshark (*Callorhincus milii*) reference genome. ANOVA KEGG pathway analysis revealed that RNA from the sharks manifested significantly higher expression than those of rays/skates in 4/5 major DNA repair pathways (Base Excision, Nucleotide Excision, Mismatch Repair, and Homologous Recombination). Each of the four pathways of DNA repair manifested differential expression of pathway-specific genes (*mpg, polL, parp4, polD2, xpa, gtf2h3, gtf2h5, ercc8, ercc4, cul4b, rad51D, rad51C, blm, ssbp1, top3a, rad51, xrcc3, mre11a, brip1, rad54b*, etc.). One gene, *polD2*, was consistently elevated in the sharks vs. rays/skates in all four pathways. A subunit of *polD*, has been proposed to contribute to the spreading and amplification of hypermutations in sharks by generating higher diversity of the T cell receptor repertoire. With increased expression of four major DNA repair pathways, sharks may be more successful than rays/skates in remediating DNA damage and surviving the genotoxic effects of increasingly polluted oceans and possibly eluding cancer.

## Introduction

Maintaining DNA integrity is critical for preventing diseases such as cancer. Assessing DNA repair and the potential for genetic damage is essential for identifying environmental threats to ocean life and informing conservation efforts. Pharmaceuticals, agrochemicals, and environmental shifts due to climate change all present threats to marine life that involve DNA damage and repair. Although saltwater studies have previously assessed crustaceans, bivalves and bony fishes (Clade *Osteichthyes*), fewer DNA damage and repair related studies have been performed on cartilaginous fishes (Clade *Chondrichthyes*), which includes Subclass Holocephali (chimaeras) and Subclass Elasmobranchii (sharks, rays, skates) (**Fig. 1**) (Alves, 2022; Consales and Marsili, 2021, Marques, 2025).

**Fig. 1.**
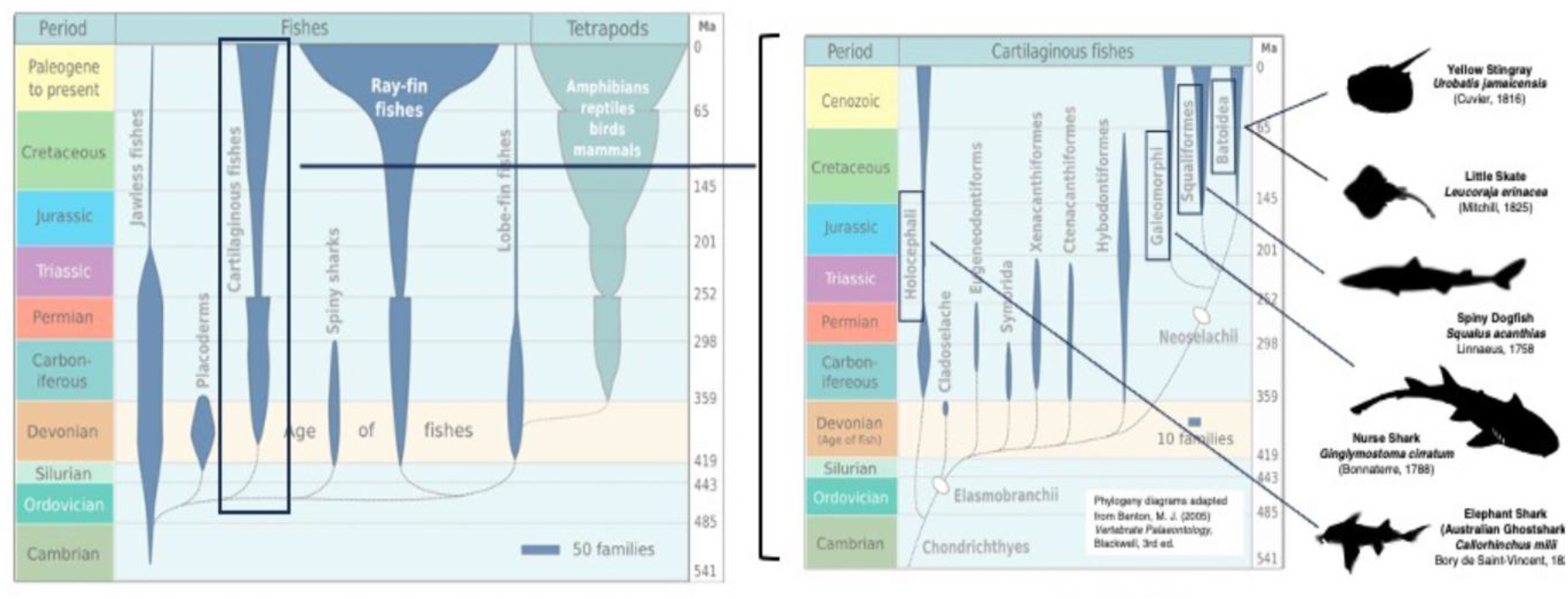
Evolution of cartilaginous fishes from the Devonian to the present to show time scales of each phylogeny divergence. The width of the spindles in each sub-figure are proportional to the number of families as a rough estimate of diversity. On the right are visual representations of each of the cartilaginous Chondrichthyan fish species used in the analysis, corresponding to different phylogenetic groups. MA indicates millions of years ago. Evolution diagrams from Epipelagic under Creative Commons license; black fish images are public domain obtained through PhyloPic.org.

All the major DNA repair pathways have been found in bony fishes within Infraclass Teleostei (Kienzler et al., 2013). Additionally, many teleosts have an additional repair pathway for UV damage known as photoenzymatic repair (*per*), which is absent in placental mammals. None of the five major DNA repair pathways have previously been demonstrated in elasmobranchs, illustrating the novelty of this study.

Elasmobranch fishes have ecological relevance because of their relatively long life-spans and broad geographic ranges, with an additional role as a food source to humans. Ecologically, elasmobranchs generally occupy high trophic roles in many marine ecosystems and exert top-down pressures in local food webs. As predators, elasmobranchs also have associated vulnerability to the bioaccumulation of chemical contaminants.

Because of overfishing and environmental contamination, over a third of elasmobranch species are threatened (Dulvy et al., 2021), especially those with coastal, sedentary lifestyles or those that are directly in contact with the oceanic sediment. Chemical contamination assessments of elasmobranchs involve accumulation of metals and organic pollutants known to cause DNA alterations and DNA damage (Alves, 2022; Chynel, 2021). Global climate change has caused variations in oceanic salinity and acidification, complications that can contribute to DNA alterations (Alva, 2017; Alves, 2022). The research on DNA damage in ocean species has primarily addressed these risks in bony fishes (Wirgin and Walman, 1998; Simoniello et al., 2009; Canedo et al., 2021).

Little is known about the pollution tolerance of elasmobranchs. Reviews like Gelsleichter & Walker (2010) and Bezerra et al., (2019) focus mostly on contaminant levels rather than DNA repair mechanisms to remediate specific forms of DNA damage. Other reviews like Tiktak et al., (2020) focus on pollutants from a human consumption perspective. Marra et al., (2019) found molecular sequences associated with DNA repair mechanisms in the great white shark genomes (*Carcharodon carcharias*) but did not assess relative RNA expression or functionality. However, many elasmobranchs spend time in coastal, likely more polluted waters for at least reproductive purposes, such as nursery grounds for pups and egg cases (Heupel et al., 2018).

The present study provides novel information on DNA repair pathways (**Fig. 2**) for four coastal elasmobranchs. Chondrichthyan fishes (combined Holocephalan chimaeras and elasmobranch sharks, skates, and rays) diverged from bony fishes – including the sarcopterygian fishes (and Tetrapoda) during the Ordovician Period (485-443 mybp [millions of years before the present]) (**Fig. 1**). The fish sampled in this study are shown in black in **Fig. 1** as well as the species used for the reference genome. Batoids are one of the more recent groups to appear in Neoselachii, appearing at the end of the Jurassic Period (145 mybp). Note that these eras are only approximate, since most fossils of elasmobranchs are from teeth, dentine-covered placoid scales, and other hard structures (e.g., dorsal spines on spiny dogfish), which makes specimen-based, physical character phylogenies difficult. However, these broad phylogenetic differences between Galeomorphs and Squaliforms are consistent throughout several analyses (Maisey et al., 2004; Naylor et al., 2012), especially as distinct from Batoidea (Aschliman et al., 2012). Others have repeatedly confirmed the differences between Rajids (skates) and Myliobatiformes (rays) (Villalobas-Segura et al., 2022).

**Fig. 2.**
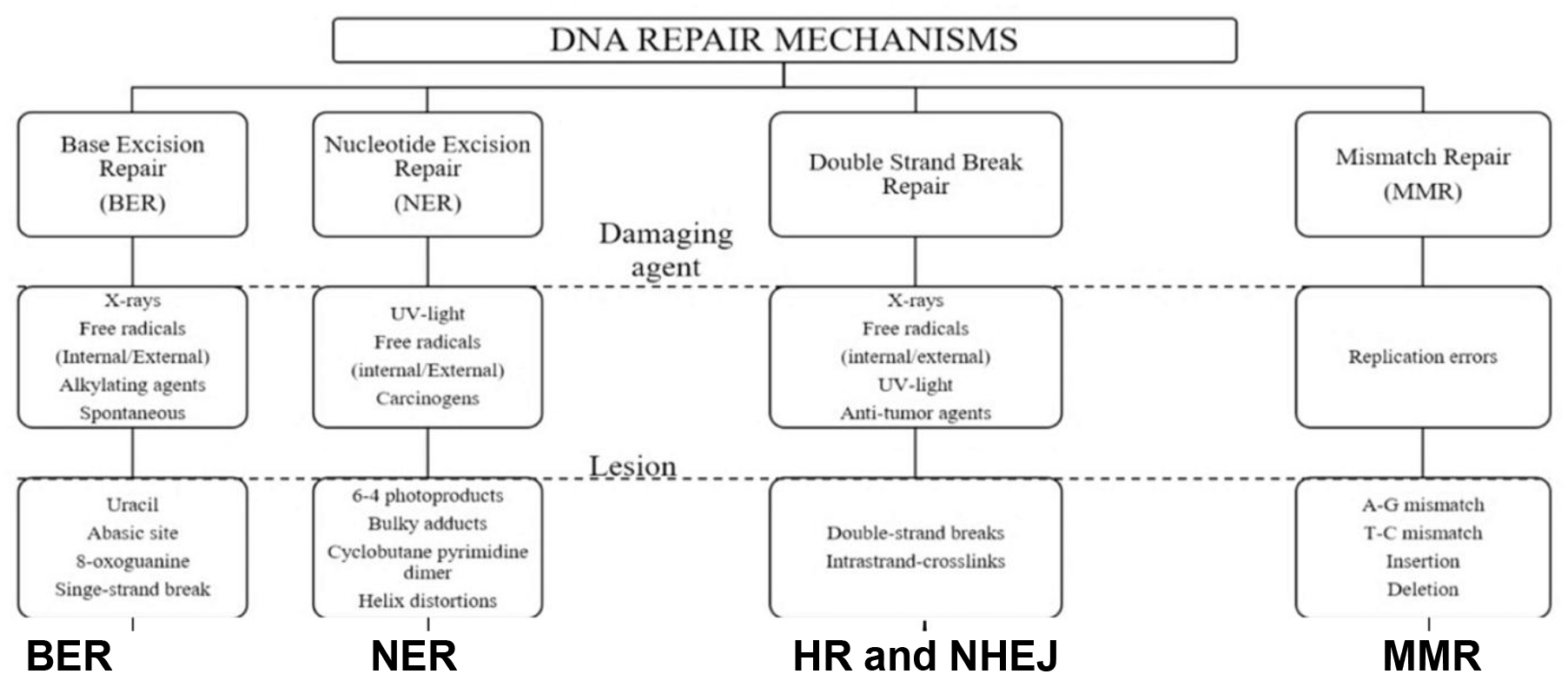
The five major pathways of DNA repair are summarized.

## Methods

### Elasmobranch sampling

All the work in this study was covered by NSU IACUC protocol 2017-DK-01 to DWK. Nurse sharks (*Ginglymostoma cirratum*, n=3) and yellow stingrays (*Urobatis jamaicensis*, n=3) were hand captured near Fort Lauderdale, Florida, while spiny dogfish (*Squalus acanthias*, n=3) and little skates (*Leucoraja erinacea*, n=3) were captured by commercial fishing vessels near Chatham, Massachusetts. Specimens were brought onto the fishing vessel or dock, and then sterile techniques were performed on the live animal to remove a small sample of skin. The tissue sample was then placed into a sterile, RNAse-free Eppendorf tube and snap-frozen in a dry ice and ethanol mixture. Samples were subsequently stored at -80°C.

### RNA Isolation and Sequencing

Tissue was pulse homogenized on ice for 20-30 seconds in the presence of RNAse inhibitor in a fume hood. RNA was isolated using the Qiagen RNAeasy Kit, and RNA integrity and concentration were determined using an Agilent TapeStation. RNA samples were then subjected to library preparation using Illumina’s TrueSeq Stranded Total RNA library generation kit and sequenced on a 2x150 base-pairs paired-end run using the NextSeq 500 High Output (300-cycle; 400 million read flow cell). The data were delivered as Fastq files, which were analyzed using Partek Flow software (Partek Inc., MO).

### Data Analysis

Alignment of exonic reads was performed using BWA-MEM (Burrows-Wheeler Aligner) in Partek Flow software using the Australian ghost shark (*Callorhincus milii*) reference genome. Aligned reads were then quantified using the Partek Annotation Model E/M to produce gene counts. Gene set ANOVA (FDR value of ≤0.05), an unsupervised form of data analysis, was used to determine statistically significant pathway differences between the sharks vs. rays/skates. Since four major DNA repair pathways were among the top 50 significant pathways, we report those here.

RNA sequencing results were based on three biologic RNA samples from each species of elasmobranch. Four species in total were used (2 sharks and 2 rays/skates) and 3 individuals per species were included. Data were expressed as the mean ± standard error for each group. An FDR value of ≤0.05 was used to determine significance in ANOVA pathway analyses for KEGG pathways which included pathways of DNA repair (**Table 1** last column).

**Table 1.** Unsupervised KEGG pathway analysis. DNA repair related pathways that show differential expression (for the entire pathway) for sharks vs. rays/skates using Gene Set ANOVA . 4/5 major DNA repair pathways were among the top 50/164) significant pathways; also included was the Fanconi Anemia pathway which overlap several major DNA repair pathways. P values and FDR (False Discovery Rate) values were evaluated for these pathways, as a whole. Pathways are ranked using the more stringent FDR step up criterion. NHEJ is the only major pathway of DNA repair that is not differentially expressed between these groups. Fanconi anemia is a disease-related pathway. It includes aspects of NER (Nucleotide Excision Repair) and of HR (Homologous Recombination).

| Gene Set Description | Fold Change | P Value | FDR Step Up |
| --- | --- | --- | --- |
| Homologous Recombination | 5.42 | 3.43E-7 | 6.24E-6 |
| Mismatch Repair | 11.08 | 4.35E-6 | 4.75E-5 |
| Nucleotide Excision Repair | 4.76 | 1.01E-6 | 1.38E-5 |
| Base Excision Repair | 8.88 | 1.41E-4 | 9.65E-4 |
| Fanconi Anemia | 1.78 | 3.13E-5 | 2.6E-4 |

Unpaired, one-tailed Student’s t tests using a p value of ≤ 0.05 were performed on individual genes in each DNA repair pathway using Microsoft Excel (version 16.63) and GraphPad (version 9.3.1). Finally, normalization was performed using counts per million reads.

## Results

There are multiple types of DNA damages (base damage, bulky adducts and single-stranded crosslinks, double stranded breaks (blunt-ended or sticky ended) and mismatched bases after replication. These damages are resolved by five distinct pathways of DNA repair (**Fig. 2**). Base Excision Repair (BER) and Nucleotide Excision Repair (NER) remediate abnormal geometry in the DNA helix generated from single strand damages. BER repairs only the base that is damaged using evolutionarily conserved and specific glycosylases. NER resolves damages involving alkylating agents (common as pollutants) and single-stranded crosslinks by removing the area around the damage and resynthesizing a long patch (27-30 bp). Double stranded helix damage is resolved either by homologous recombination (HR) or non-homologous end joining (NHEJ). HR uses a sister chromatid as the template for slow, high fidelity repair, whereas NHEJ is rapid and error prone in comparison.

An examination of the top 50 out of a total of 164 significant KEGG pathways differentially expressed between sharks vs ray/skates included four of the five major DNA repair pathways (**Table 1**). These include BER (**Fig. 2, Table 1** and individual genes shown in **Fig. 3**), NER (**Fig. 2, Table 1** and individual genes shown in **Fig. 4**), HR (**Fig. 2, Table 1** and individual genes shown in **Fig. 5**) and MMR (**Fig. 2, Table 1** and individual genes shown in **Fig. 6**) using Gene Set ANOVA analyses. The Fanconi Anemia pathway was among the top 50 (**Table 1**, and individual genes shown in **Fig. 7**) and is shown here because it is a disease state that includes components of double stranded and single stranded DNA repair pathways. It also includes related pathways such as cell cycle. The only DNA repair pathway not shown to be differentially regulated by sharks vs. rays/ skates was NHEJ. Given the critical importance of DNA repair mechanisms in the cell, it is remarkable that 4/5 pathways of repair show the same trend (higher in sharks).

**Fig. 3.**
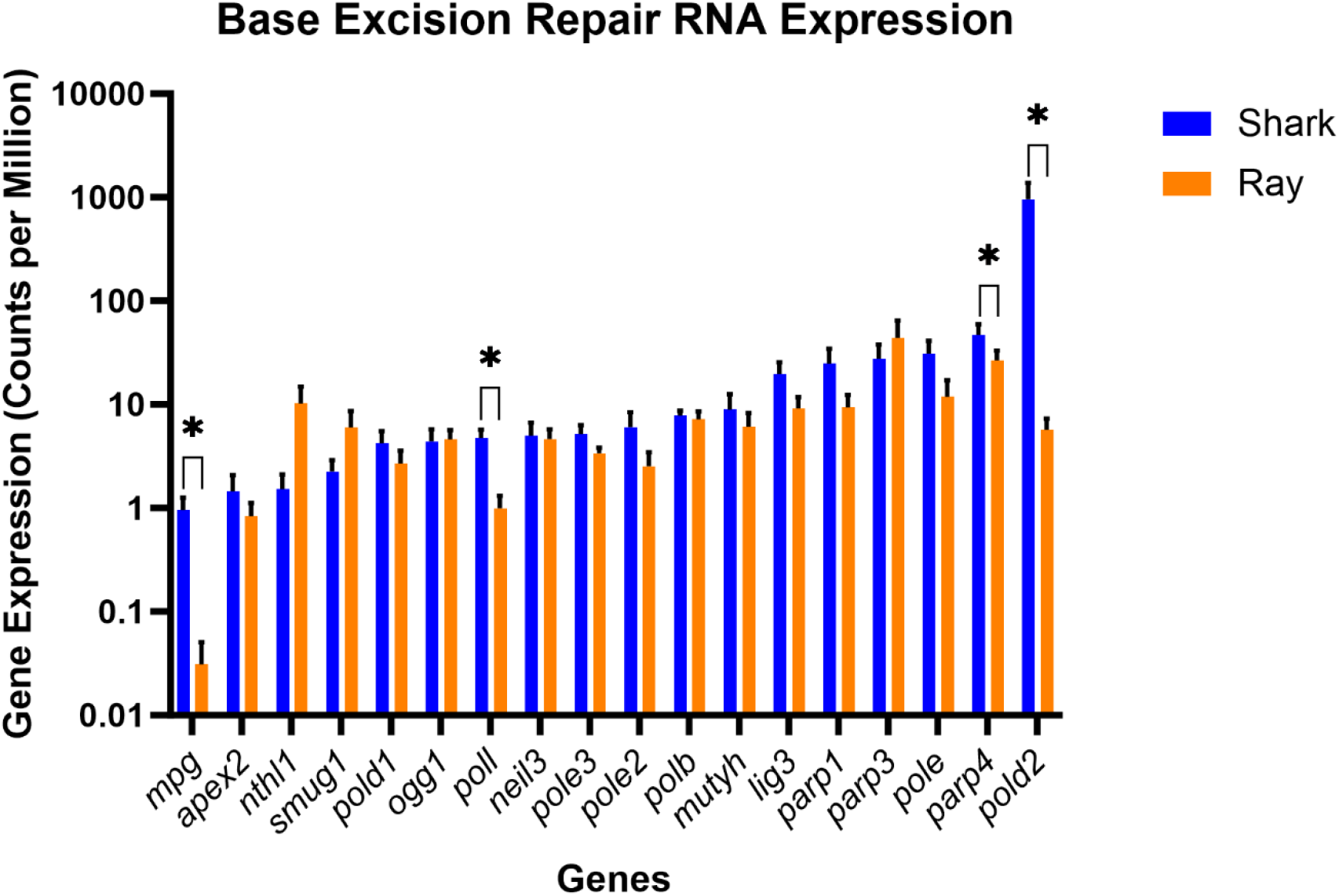
RNA expression via RNA sequencing for genes involved in the **Base Excision Repair Pathway**. Significantly different steady state RNA expression was indicated with an asterisk (*) p≤ 0.05. 4/18 genes were significantly higher in sharks vs. rays/skates: *mpg, polL, parp4* and *polD2. polD2* is the major polymerase involved in replication and repair.

**Fig. 4.**
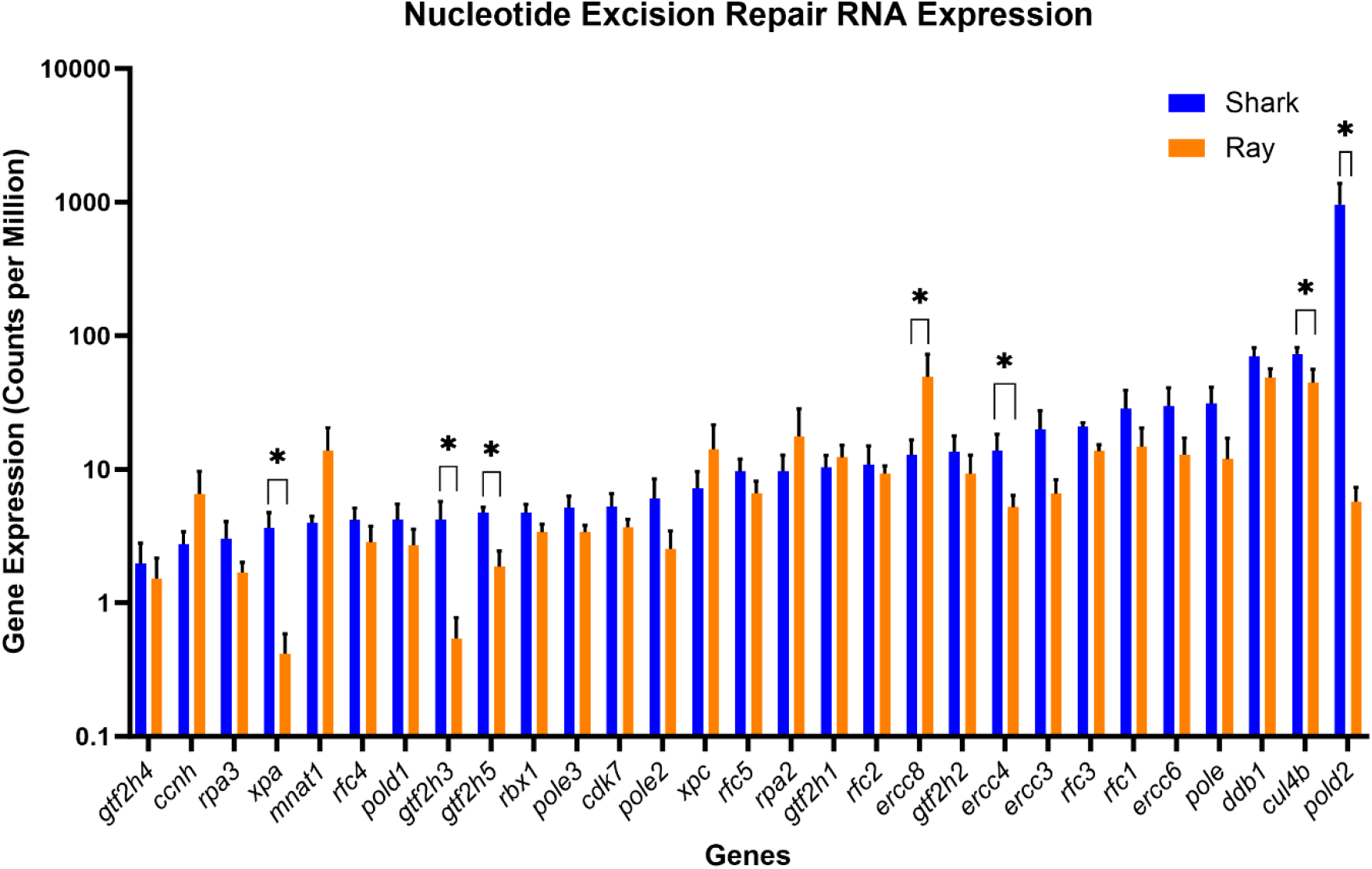
RNA expression for genes involved in the **Nucleotide Excision Repair Pathway**. Significantly different steady state RNA expression was indicated with an asterisk (*) p≤ 0.05. 6/29 genes were significantly higher in sharks vs. rays/skates: *xpa, gtf2h3, gtf2h5, ercc8, ercc4, cul4b*, and *polD2*. 1/29 genes was significantly higher in rays/skates compared with sharks: *csa/ercc8* which is involved in transcription coupled NER.

**Fig. 5.**
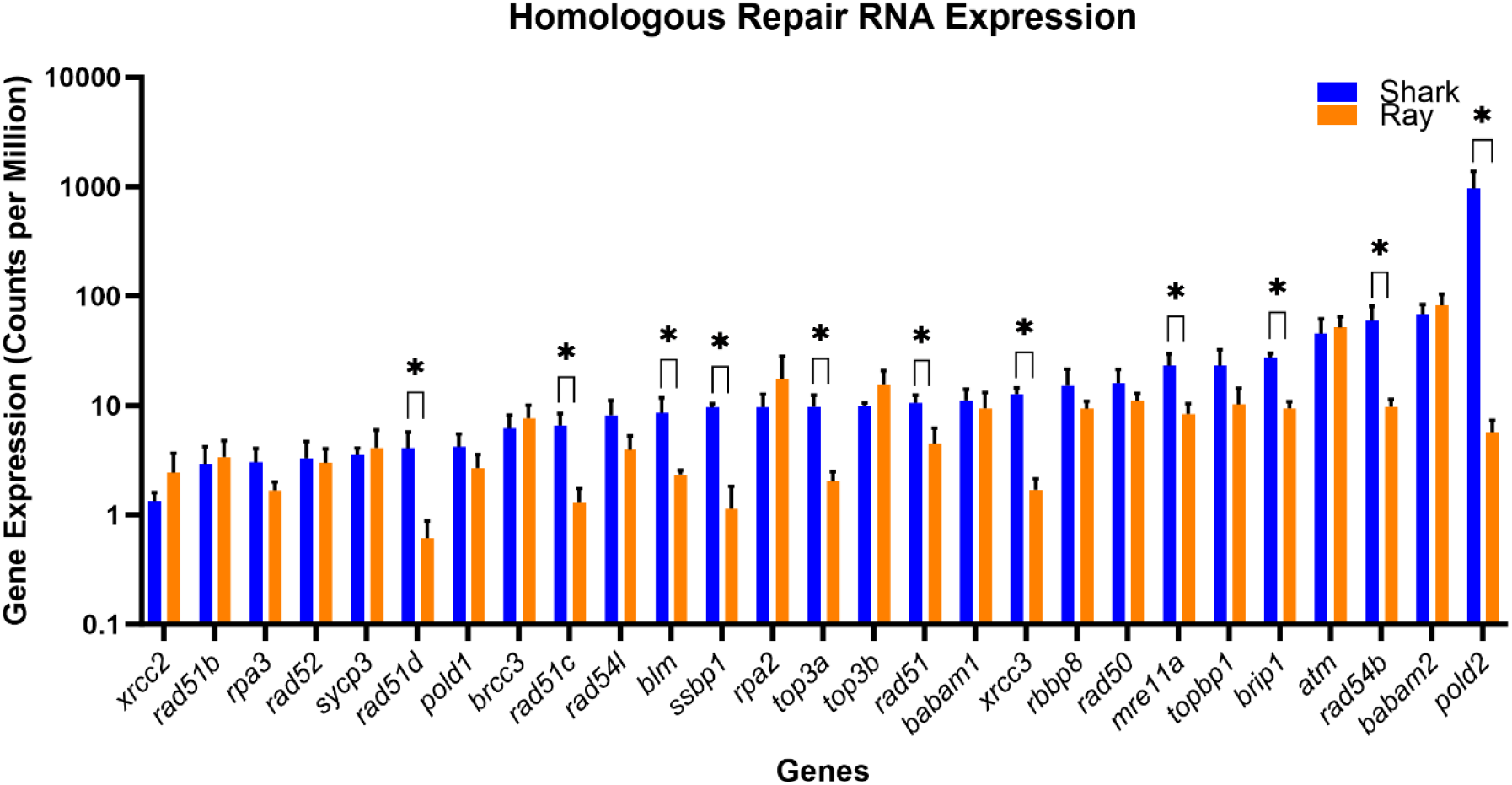
RNA expression for genes involved in the **Homologous Recombination Repair Pathway**. Significantly different steady state RNA expression was indicated with an asterisk (*) p≤ 0.05. 11/27 genes show significantly higher shark vs. ray/skate expression: *rad51D, rad51C, blm, ssbp1, top3a, rad51, xrcc3, mre11a, brip1, rad54b*, and *pold2*.

**Fig. 6.**
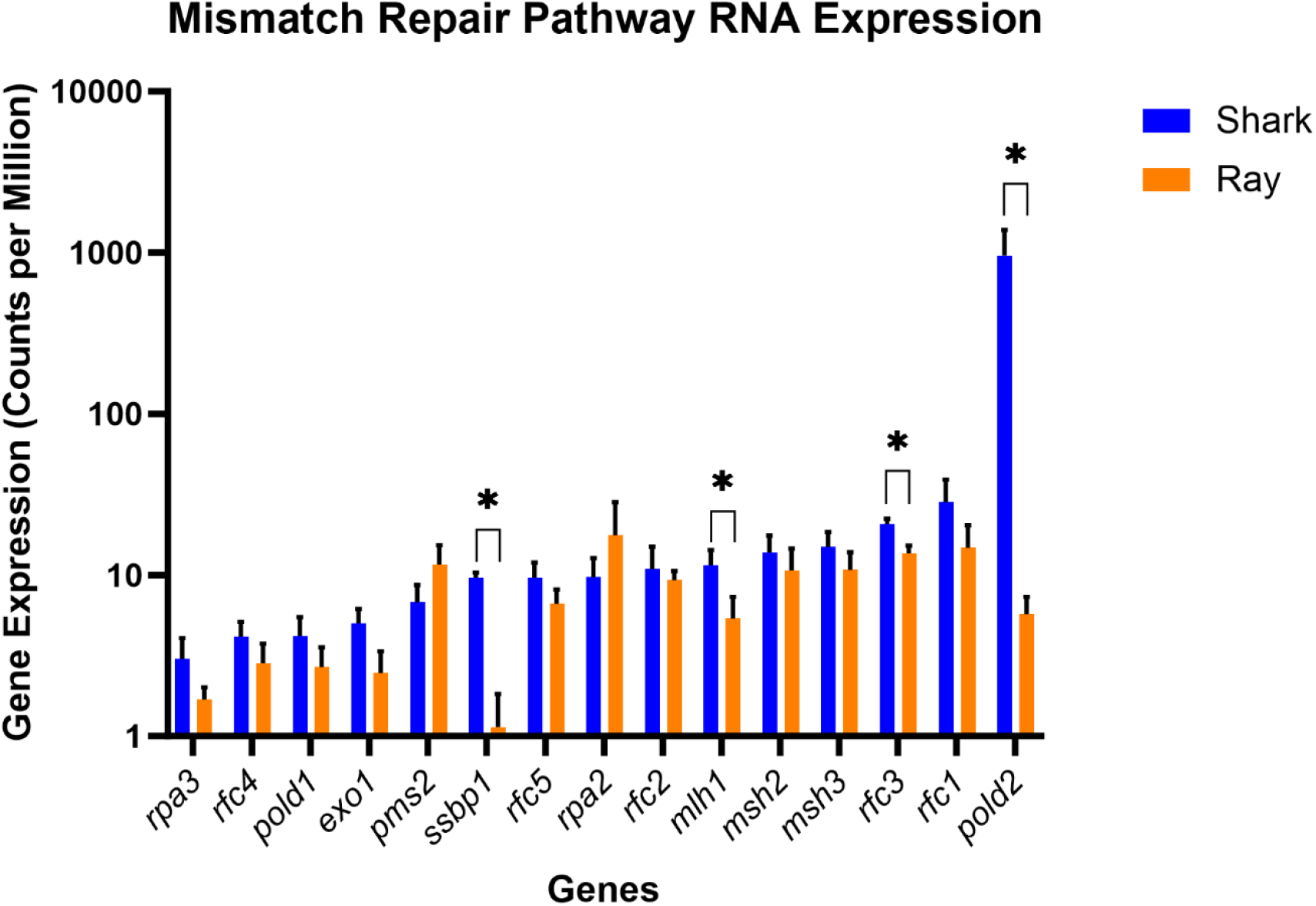
RNA expression for genes involved in the **Mismatch Repair Pathway** which is a post-replicative repair pathway. Significantly different steady state RNA expression was indicated with an asterisk (*) p≤ 0.05. 4/15 genes show significantly higher RNA expression in sharks vs. rays/ skates: *ssbp1, mlh1, rfc3*, and *polD2*.

**Fig. 7.**
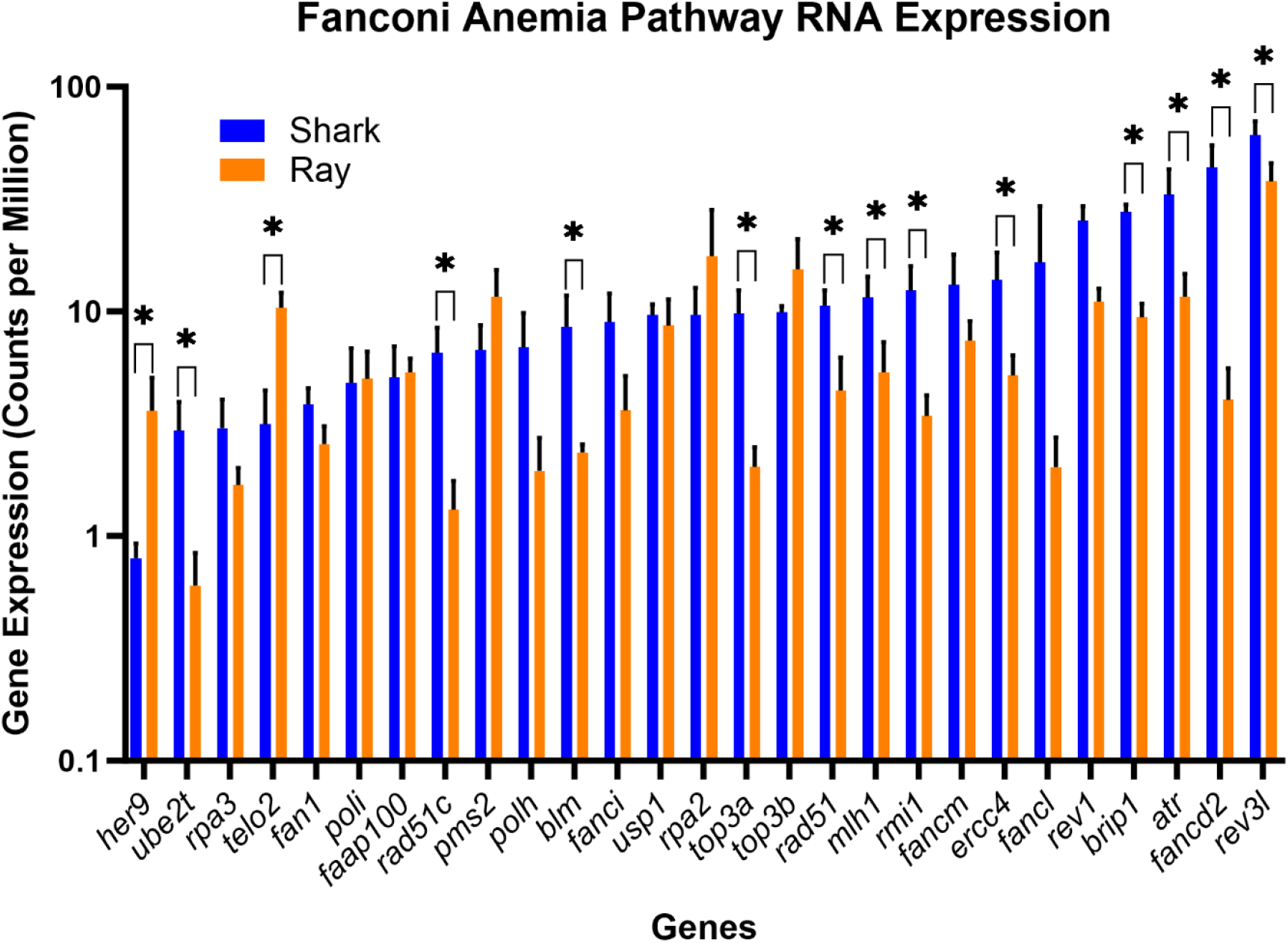
RNA expression for genes involved in the **Fanconi Anemia Pathway**. Significantly different steady state RNA expression was indicated with an asterisk (*) p≤ 0.05. 12/27 genes show significantly higher RNA expression in sharks vs. rays/skates: *ube2T, rad51C, blm, top3A, rad51, mlh1, rmi1, ercc4, brip1, atr, fancd2, rev3l*. Two genes (*her9, telo2*) show the opposite pattern with sharks being lower than rays/skates.

We subsequently performed an analysis of pathway specific genes using unpaired Student’s t tests identifying the genes that differentially regulated (p ≤ 0.05)(**Figs. 3-7**). In addition, genes common to multiple pathways of DNA repair were identified (*ercc4, rad51, pold2*). The polymerase delta2 subunit of *polD* was common to multiple pathways (BER, NER, HR and MMR) and significantly higher in sharks vs. rays/ skates. Polymerase delta is indispensable for leading and lagging strand synthesis in replication. *polD2* does not encode the catalytic subunit.

There were also pathway specific genes in each of the four pathways showing higher expression in sharks vs. ray/skates, so the pathway statistical trends identified in the ANOVA analysis were not driven by a small number of overlapping genes.

Base Excision Repair is a pre-replicative repair pathway that repairs small base lesions caused by alkylation, oxidation and deamination. For Base Excision Repair four genes were significantly higher in sharks vs. rays/skates: *mpg, polL parp4*, and *polD2. polD* is the major polymerase involved in replication and repair. N-methylpurine-DNA glycosylase (*mpg*) initiates BER by removing a large variety of deaminated, alkylated and lipid-peroxidation induced purine adducts. At least 11 distinct glycosylases are known in BER in mammals (Krokan and Bjoras, 2013).

NER is a pre-replicative DNA repair pathway which causes the replication fork to pause so that the repairosome can remediate the damage before DNA is replicated, avoiding mutation in the progeny strands. Six genes were significantly higher in sharks vs. rays/skates: *xpa, gtf2h3, gtf2h5, ercc4, cul4b*, and *polD2*. The *gtf2h* gene products are involved in DNA unwinding, recruitment of downstream DNA repair factors and verification of bulky lesions on DNA. These proteins are part of the transcriptional machinery as well as transcription coupled NER (Wu et al., 2008). *xpa* in concert with damaged DNA initiates repair by binding to damaged sites and stimulates the 5’-3’ helicase activity of *xpd* and the DNA translocase activity of *xpb* (Fadda, 2015). The product of the *ercc4/xpf* gene interacts with *ercc1* which plays a crucial role in DNA repair and recombination (Krasikova et al., 2021). The *ercc8/csa* gene product is involved in transcription-coupled NER as opposed to global genomic NER (Saijo, 2013). This gene is expressed at higher RNA levels in rays/skates vs. sharks.

The pathway of DNA repair that did not show differential RNA expression between these groups was NHEJ (data not shown). NHEJ exists in zebrafish so the finding that NHEJ is not differentially expressed in elasmobranchs cannot be explained by lack of homology. NHEJ and HR both repair double stranded breaks, however NHEJ is a faster, more efficient pathway directly ligating the broken ends of DNA in double strand breaks, although often with some loss of information. NHEJ is inherently more mutation generating than HR (Stinson and Loparo, 2021).

In the HR pathway eleven genes showed significantly greater expression in sharks vs. rays/skates (*rad51d, rad51c, blm, ssbp1, top3A, rad51 xrcc3, mre11a, brip1, rad54b, polD2*). *rad51c* and *rad51d* are tumor suppressor genes in mammals with known mutations associated with breast and ovarian cancers. The *xrcc3* gene product, a tumor suppressor gene, is part of the *rad51* paralog protein complex (Uhrig et al., 2024). The *top3A* gene product (topoisomerase 3 alpha) resolves Holliday junctions (replication forks) and is implicated in Bloom’s syndrome-like disorder when mutated (Martin et al., 2018).

Mismatch Repair is a post replicative repair process. In Mismatch Repair, four genes show significantly higher expression in sharks vs. rays/skates (*ssbp1, mlh1, rfc3, polD2*). The replication protein C factor (*rfc*) plays an important role in MMR, acting as a clamp loader, loading *pcna* onto DNA which is essential for the *mut* endonuclease activity on the mismatched nucleotide (Skibbens et al., 2005). *mlh1* is part of a protein complex that detects and binds to mismatches. Mutation in this gene is associated with colorectal cancer (Pećina-Šlaus et al., 2020). *mlh1* promoter methylation is now considered to be the primary epigenetic cause of MMR deficiency, leading to microsatellite instability-high sporadic cancers (Rico-Mendes et al., 2025). Genes that overlap in two or more pathways include *polD2, rfc, ercc4, Rad51*, and *ssbp1*.

The Fanconi Anemia (FA) pathway is included in this analysis although it is not a pathway of DNA repair but is instead a disease state, that includes aspects of double strand and single strand repair as well as replication (Moreno et al., 2021). In addition, the FA pathway also reveals functions that are related to other aspects of genomic instability like cell cycle regulation. The *atr* gene encodes a protein kinase, a monitor of DNA damage that is involved in cell cycle checkpoint regulation upstream of DNA repair pathways. The 12 genes expressed at higher levels in shark vs. ray/skate are: *ube2T, rad51C, blm, top3A, rad51, mlh1, rmi1, ercc4, brip1, atr, fancd2, rev3L*. The gene *ercc4* is part of the NER pathway. The *mlh1* gene product is part of mismatch repair. The gene product of *top3A* (topoisomerase gene) relieves supercoiling upstream of replication forks and is included in the replication pathway. The *rad51* and *rad51c* genes are part of the HR pathway (Uhrig et al., 2024). The *brip1 gene* encodes a DNA helicase that interacts with the *brca1* gene (Wang et al., 2023).

## Discussion

The study of wild species that reflect the unusual evolutionary success represented by the shark is important for our understanding of fundamental biological processes like DNA repair. The implications for the avoidance of cancer in increasingly polluted oceans are also important from the dual perspectives of increased immunity and DNA repair.

Numerous freshwater fishes have been studied for functional DNA repair (reviewed by Keinzler, et al., 2013), including rainbow trout (*Oncorhynchus mykis*s) and zebrafish (*Danio rerio*). However, similar DNA repair studies are presently lacking for many marine species, including marine elasmobranchs. Fewer still have involved examination of the transcriptome for DNA repair pathways as a whole or assessing the expression of individual genes that are unique to these pathways and are involved in the genesis of cancer. In addition to pathway analyses, our study has shown that the gene *polD2* which is differentially expressed in all of these pathways, is higher in sharks than rays/skates.

Despite a close phylogenetic relationship between the four species of elasmobranchs, we see a clear distinction in gene expression patterns for 4/5 major pathways of DNA repair with the two sharks being significantly higher in gene expression. The related pathways of DNA replication and p53 signaling show the same trend (data not shown). Although DNA repair and replication are related, one does not drive the other; in other words, high replication does not necessarily correlate with high repair (Latimer et al., 1996; Latimer et al., 2003).

Most DNA repair genes are tumor suppressor genes. Many of the genes that show differential gene expression in this study with higher expression in sharks vs. rays/skates are mutated in human cancers (e.g. *brip1* (Suszynska et al 2020), *top3a* (Broberg et al 2009), *xpa* (Feltes and Bonatto, 2015), *rad51, atr, mlh1* (Song et al 2015), *ercc4* (Bogliolo et al 2013), *polD* (Gola et al 2023; Mendiratta et al., 2021). *mlh1* has particular importance as an epigenetic regulator of microsatellite instability (when downregulated) and is used as a biomarker to determine if a cancer will be high mutational burden and therefore a candidate for immunotherapy (Rico-Mendez et al., 2025). One could speculate that the expression of these genes may be correlated with a relative lower rate of cancers in sharks (Sendell-Price et al., 2023) since they are all expressed at higher steady stage RNA levels in the sharks vs rays/skates. With greater study of wild species this may become apparent.

In our previous work in cancer, we have found a strong correlation between the transcriptomic patterns, protein expression and functional repair for NER (Latimer et al., 2010) particularly when assessed for the entire pathway of repair. NER enzymes are expressed at low levels compared to many other classes of genes (Christmann and Kaina, 2013). Rather than finding a few genes differentially expressed we have found 4/5 DNA repair pathways as a whole, are in the top 50 most significantly regulated pathways out of 164. The remarkable findings of the present study suggest that sharks could be more successful in surviving polluted waters containing genotoxic contaminants than rays/skates based on 4/5 significantly different pathways of DNA repair (Everaarts and Sarkar, 1996). One can envision these repair pathways being advantageous in the context of oil spills, for example. Carcinogens (specifically alkylating agents) are present in oil and petrochemicals (Rodrigues et al., 2010). Chemicals used to disperse oil, such as Corexit 9527A which contains 2-Butoxyethanol, permeabilize cell membranes thus creating greater nuclear exposure from these genotoxic carcinogens (Hoflack, 1997).

An alternative aspect of DNA repair involves somatic hypermutation which has been found in sharks in T cell specific genes *tcr* gamma and delta variable regions (Diaz et al., 1998; Zhu and Hsu, 2010). This hypermutation is believed to play a role in increasing the diversity of the T cell receptor repertoire, allowing sharks to respond to a wider variety of pathogens. This hypermutation is a function of certain DNA repair related genes such as error prone DNA polymerases (E, epsilon or D, delta) (Diaz et al., 1998; Zhu and Hu, 2010). Another potential ramification of increased T cell receptor diversity and increased DNA repair in general, is the ability to elude cancer.

Although hypermutation is documented in sharks in cells of the immune system, this study used the skin of all four species of elasmobranchs. Immune cells could have been present in the skin. NER is known to be tissue specific in mammals (Latimer et al.,1996; Latimer et al., 2003; Latimer et al., 2008; Latimer et al., 2015) and remediates two types of UV based damages. HR is also known to be tissue specific in mammals, being more active in stem cells and actively proliferating cells (Tichy et al., 2010; Bishop and Schiestl, 2002). Tissue specific DNA repair has not yet been explored in elasmobranchs but could be a opportunity for future research, particularly between tissues of sensitivity (e.g., gill filaments) or known sites of toxin accumulation (e.g., liver).

Sharks have existed on this planet 500 million years whereas *H. sapiens* have existed 300,000 years. The study of sharks and other elasmobranchs is essential for our understanding of biochemical processes related to genomic stability like DNA repair.

## Acknowledgements

Funding was provided by the NSU President’s Faculty Research Grant Program (award to JJL and DWK) and the AutoNation Institute for Breast Cancer Research and Care (JJL). We appreciate the vessel time by Captains Doug Feeney (F/V *Noah*) and Dave Gelfman (F/V *Horse Mackerel*), plus the field assistance by Jacqueline Reuder (NSU). Many thanks to Stephen Zimberg and Carol Murphy for their critical reading of this work. Special acknowledgement goes to Jennifer Kalil for her work on figure preparation. This paper is dedicated to the memory of Professor Arnold Miller for instilling a love of science in all who knew him.

